# Base-rate neglect and neural computations for subjective weight in probabilistic inference

**DOI:** 10.1101/671396

**Authors:** Yun-Yen Yang, Shih-Wei Wu

## Abstract

Humans show systematic biases when estimating probability of uncertain events. Base-rate neglect is a well-known bias that describes the tendency to underweight information from the past relative to the present. In this study, we characterized base-rate neglect at the computational and neural implementation levels. At the computational level, we established that base-rate neglect arises from insufficient adjustment to weighting prior information in response to changes in prior variability. At the neural implementation level, we found that orbitofrontal cortex (OFC) and medial prefrontal cortex (mPFC) represent subjective weighting of information that reflects base-rate neglect. Critically, both subjective-weight and subjective-value signals that guide choice were found in mPFC. However, subjective-weight signals preceded subjective-value signals. These results indicate that when facing multiple sources of information, estimation bias such as base-rate neglect arises from information weighting computed in OFC and mPFC, which directly contributes to subjective-value computations that guide decisions under uncertainty.

**Significance Statement:** Facing uncertainty, estimating the probability of different potential outcomes carries significant weight in affecting how we act and decide. Decades of research show that humans are prone to giving biased estimation but it remains elusive how these biases arise in the brain. We focus on base-rate neglect, a well-known bias in probability estimation and find that it is tightly associated with activity in the medial prefrontal cortex and orbitofrontal cortex. These regions represent the degree to which human participants weigh different sources of information, suggesting that base-rate neglect arises from information-weighting computations in the brain. As technology provides us the opportunity to seek and gather information at an ever-increasing pace, understanding information-weighting and its biases also carry important policy implications.

## Introduction

Base-rate neglect – the tendency to underweight base rate or prior information compared with current, individuating information – is a well-known bias in human probabilistic inference (Tversky & Kahneman, 1973, 1974). It highlights a long-lasting research program in experimental psychology that uses the ideal Bayesian inference as a model for human performance and seeks to gain insights into the nature of human inference by identifying systematic deviations in actual performance from the ideal model prediction (Edwards, Lindman, & Savage, 1963). In the case of base-rate neglect, despite its popularity, its causes, ecological validity and the degree of base-rate underweighting remain debated (for review, see Koehler (1996)).

In cognitive neuroscience, there is a growing interest in studying the neurocomputational substrates involved in a variety of inference tasks, from how people use cue reliability as prior information to guide perceptual decision making (Forstmann, Brown, Dutilh, Neumann, & Wagenmakers, 2010; Mulder, Wagenmakers, Ratcliff, Boekel, & Forstmann, 2012), make financial decisions based on prior information about partner’s reputation (Fouragnan et al., 2013), combine prior and likelihood information about reward probability (d’Acremont, Schultz, & Bossaerts, 2013; Ting, Yu, Maloney, & Wu, 2015), to infer latent causes (Chan, Niv, & Norman, 2016) and other people’s intentions (Chambon et al., 2017). These studies pointed to the role of medial prefrontal cortex (mPFC) and orbitofrontal cortex (OFC) in inference, from representing prior information (Forstmann et al., 2010; Fouragnan et al., 2013; Ting et al., 2015; Vilares, Howard, Fernandes, Gottfried, & Kording, 2012), current observation or sensory evidence (d’Acremont et al., 2013; Ting et al., 2015) to the combination of prior and current information (Chambon et al., 2017; Chan et al., 2016; Ting et al., 2015).

An important but open question is what are the neural mechanisms contributing to the systematic biases in inference found in the behavioral literature. In this study we seek to address this question by focusing on base-rate neglect. However, there are at least three major challenges in addressing this question. First, in task paradigms suitable for neurobiological investigations, the field has yet to see paradigm(s) that robustly reveals or replicates base-rate neglect. Across the studies highlighted above, subjects either achieved near-optimal performance in combining prior and current information (Chan et al., 2016; Ting et al., 2015) or it is unclear whether subjects achieved near-optimal performance (Chambon et al., 2017; d’Acremont et al., 2013; Forstmann et al., 2010; Fouragnan et al., 2013; Mulder et al., 2012; Vilares et al., 2012). To address this challenge, it is critical to identify the statistical properties of prior and likelihood information such that, when manipulated, produce robust base-rate neglect.

Second, there is little consensus on which behavioral metric one should use to quantitatively characterize base-rate neglect and to identify brain regions involved in computing this metric. To address this challenge, we proposed that subjective weight – how subjects weigh prior and likelihood information – should be the behavioral metric to characterize base-rate neglect, since it is fundamentally about underweighting of prior information. But how should one go about investigating subjective weight and what would be the starting neural hypothesis? Bayesian decision theory provides a key insight. That is, subjective weight assigned to prior and likelihood should be determined by the relative variability of these two sources of information. Therefore, a reasonable starting hypothesis should be that in computing subjective weight, the brain takes into account prior and likelihood variability. To test this hypothesis, it is important to independently manipulate the variability of both prior and likelihood so that we could observe how subjective weight changes in response to information variability and compare it with the ideal weight from the Bayesian integrator. At the neural implementation level, the link between information variability and subjective weight also leads to an important hypothesis. That is, neural substrates for subjective-weight computations should be tightly linked to regions that represent information variability. Together, these observations point to three critical analyses that aim to (1) identify brain activity representing subjective weight, (2) identify neural representations of information variability and (3) characterize the connecting properties between neural substrates identified in (1) and (2) at the time of prior-likelihood integration.

Finally, the third challenge is concerned with the tight relation between inference and decision making. Since the purpose of most inferences is to help guide decision-making where OFC and mPFC are also heavily involved (Bartra, McGuire, & Kable, 2013; Clithero & Rangel, 2013; Kable & Glimcher, 2009; Padoa-Schioppa & Conen, 2017), it remains unclear whether inference-related computations are dissociable from choice-related computations. To address this challenge, in our decision task we temporally separated inference from choice such that, during inference, it is not possible to engage in decision computations. Given this design, we aimed to identify subjective-weight representations during the inference stage and subjective-value representations during the choice stage of the task.

We found that subjects showed robust base-rate neglect that can be attributed to lack of sensitivity in response to changes in variability of prior information. Using BOLD fMRI, we identified neurocomputational substrates for subjective weighting of information that reflects base-rate neglect and established that subjective-weight signals were temporally dissociable from subjective-value signals that guide choice. These results indicated that estimation bias – pervasively observed in many human probability judgments – arises from information-weighting computations that are differentiable from decision computations.

## Results

Our goal was to investigate neural computations for probabilistic inference – the potential integration of prior and likelihood information. The experiment consisted of two sessions – prior-learning (Session 1, behavioral) and prior-likelihood integration (Session 2, fMRI) – run on two consecutive days. In the prior-learning session, subjects learned through feedback the probability distribution on probability of reward associated with two visual symbols, each representing a unique distribution with the same mean (0.5) but differed in variance (Fig.1A). On each trial, subjects were presented with one symbol and asked to estimate the probability of receiving a reward that was randomly drawn from the probability distribution associated with the symbol (Fig. 1B). Subjects were given feedback and rewarded based on how close his or her estimate was to the true probability of reward on that trial – an incentive compatible procedure developed to motivate subjects to learn probability distributions well (see Methods for details).

**Figure 1.**
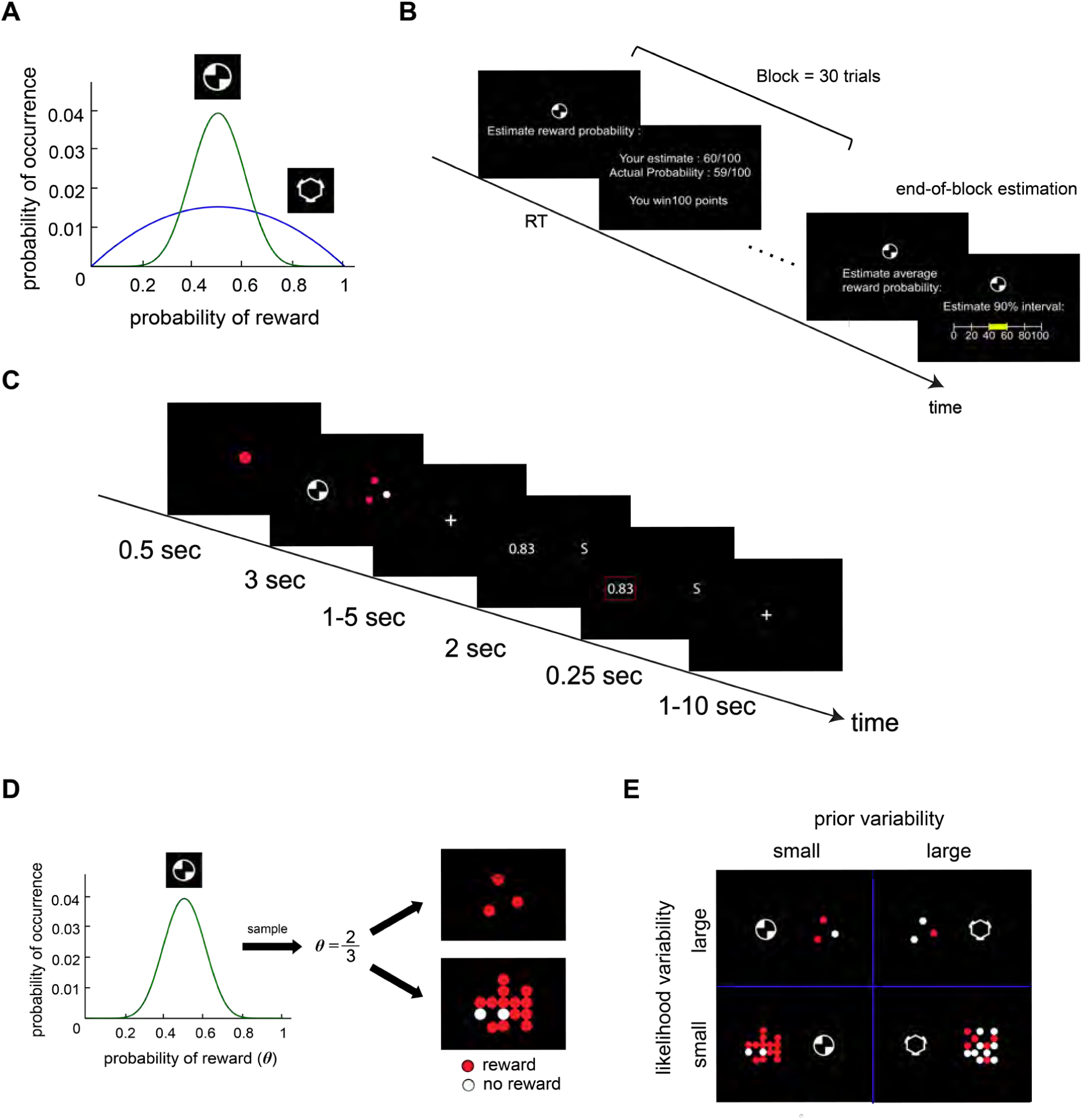
Experimental design. **A-B**. Prior-learning session. **A**. Two probability distributions on probability of reward that serve as prior information. Each distribution was represented by an abstract visual symbol. The two distributions had the same mean but different variance. **B.** Prior-learning task. Subjects learned the two distributions through feedback. On each trial, subjects were presented with a symbol and were asked to estimate the probability of receiving a monetary reward. After estimation, subjects received feedback and were rewarded based on how close his or her estimate was to the true probability of reward. **C-E**. Prior-likelihood integration session. **C**. Lottery decision task. On each trial, subjects had to choose between the symbol lottery and the alternative lottery. Prior (abstract visual symbol) and likelihood information (colored dots) associated with the symbol lottery were first presented for 3 seconds. After a variable delay, reward probability associated with the alternative lottery was revealed explicitly in numeric form. Subjects had to indicate his or her decision within 2 seconds between the symbol lottery (represented by “S” on the right side of the screen in this example) and the alternative lottery (0.83 probability of winning a reward in this example) with a button press. Once a button was pressed, subjects received a brief feedback on the chosen option (250ms). **D**. Illustrations on how likelihood information is generated on each trial. In this example, the probability of reward associated with the symbol lottery was 2/3. Likelihood information is the outcomes after repeatedly realizing this lottery – 2/3 chance of winning a reward – 3 times (screen containing 3 dots on the upper right) or 15 times (screen containing 15 dots on the bottom right). A red dot indicates a reward outcome and a white dot indicates a no-reward outcome. E. Manipulation of prior and likelihood variability. There were two levels of prior variability and two levels of likelihood variability contributing to a 2 ×2 factorial design.

One day after the prior-learning session, in the prior-likelihood integration session – an fMRI session and the primary interest of this study – subjects performed a lottery decision task. On each trial, subjects were asked to choose between two lotteries that differed only in the probability of winning a fixed monetary reward (Fig. 1C). The trial consisted of two stages: an inference stage followed by a choice stage. The goal of designing these two stages was to dissociate probabilistic inference from choice – two important computations that can be highly correlated – so that in the inference stage it is not possible for subjects to engage in decision-related computations.

In the inference stage, information about one of the lottery options, termed the symbol lottery, was presented. Two pieces of information – prior and likelihood – about the symbol lottery were shown. The prior information was one of the symbols subjects learned in the prior-learning session. The presented symbol served to inform the subjects, on the current trial, which distribution the probability of reward associated with the symbol lottery was drawn from. The likelihood information showed outcomes of repeated realizations of the symbol lottery (red dots indicated a reward outcome, white dots indicated no-reward outcome) such that if the sample size were infinitely large, the proportion of red dots would be equivalent to the probability of reward associated with the symbol lottery on the trial. Hence, when the sample size (the number of times the lottery was executed) was small, likelihood information was unreliable in indicating probability of reward. As sample size increases, the reliability of likelihood information increased (Fig. 1D). Hence, by manipulating sample size we achieved to manipulate variability of the likelihood information. In summary, we independently manipulated the variability of prior and variability of likelihood information about the symbol lottery with a 2 ×2 factorial design (Fig. 1E).

In the choice stage, information about the other lottery option, referred to as the alternative lottery, was presented. In the example shown in Fig. 1C, the alternative lottery has an 83 % chance of winning a reward. Because reward magnitude was the same between the two options, subjects should choose the option she or he thinks has the larger probability of reward. By design, the reward probability of the alternative lottery was randomly selected so that it was not possible for the subjects to predict the alternative lottery on a trial-by-trial basis. Hence, brain activity identified to be associated with probabilistic inference in the inference stage cannot be attributed to decision-related computations and brain activity identified to be associated with making a decision about which option to choose in the choice stage also cannot be attributed to inference-related computations.

### Subjects learned mean and variance of prior distributions through sampling from these distributions

We analyzed subjects’ trial-by-trial estimate on probability of reward in the prior-learning session to examine how well subjects learned the two prior distributions on probability of reward. Subjects’ probability estimates (Fig. 2A, histogram in gray) capture well the shape of the prior distributions (blue curve): the mode of both distributions was close to 0.5, the estimates were symmetric around 0.5 and were more variable when the distribution had larger variance. In addition, subjects’ estimate on the variability of reward probability associated with each symbol – provided at the end of each block of trials – also did not differ significantly from the true variability of the distributions (Fig. 2B), suggesting that they acquired accurate knowledge about the variability of prior distributions. For the probability estimate, subjects’ mean estimate did not differ from the true mean (50 % chance of reward) when prior variability was small, but was significantly smaller than the true mean when prior variability was large (t=−2.14, df=27, p=0.0416). This could be due to increased task difficulty in estimating probability under large variability (Griffin & Tversky, 1992). Such difference, however, did not change the conclusion of subjects’ behavioral performance in the subsequent session (Prior-likelihood integration session) in how they weight prior and likelihood information.

**Figure 2.**
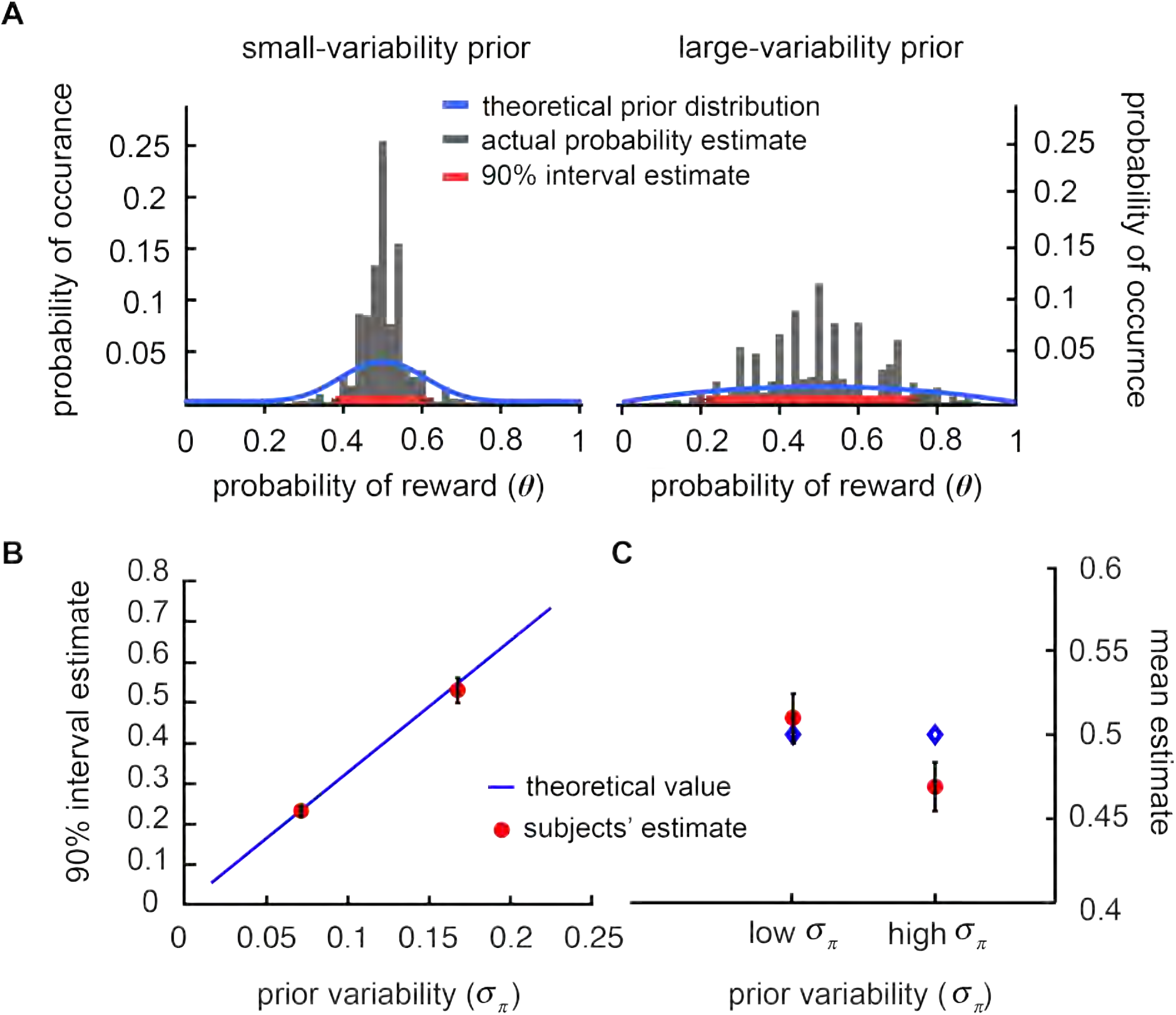
Behavioral results: Prior-learning session. **A**. Comparison of probability estimates and prior distributions. Data from all subjects’ trial-by-trial estimates on the probability of reward (*θ*) associated with the two prior distributions are summarized by the histograms in gray. The blue curves represent the prior distributions. Theta on the x-axis denotes probability of reward. **B.** Variability estimate. Subjects’ mean estimate on the 90 % interval of reward probability (data points in red) associated prior distributions is plotted against the standard deviation of the prior distributions (*σ*_π_). The blue curve represents the 90 % interval of prior distribution with standard deviation spanning from small (close to 0) to large (0.25). **C**. Subjects’ mean probability estimates (data points in red) and the true mean of the prior distributions (50 % chance of reward, in blue). Error bars represent ±1 S.E.M.

### Suboptimal integration: Subjects underweight prior information about probability of reward

In the prior-likelihood integration session, subjects were simultaneously presented with both prior and likelihood information (Fig. 1C) about the symbol lottery and had to make a choice between the symbol lottery and the alternative lottery. The choices subjects made allowed us to examine how and how well subjects combined prior and likelihood information by comparing actual performance with ideal Bayesian integration. We approached this question by estimating how the subjects weight likelihood relative to prior – referred to as subjective weight 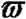 – and compared it with an ideal decision maker. To estimate subjective 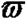, for each subject and each condition (combination of prior and likelihood variability) separately, we performed a logistic regression analysis on choice (see Behavioral analysis 1: Estimating subjective weight in Methods). If subjects completely ignored likelihood information, subjective 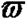 would be 0. By contrast, if subjects only considered likelihood information, subjective 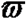 would be 1.

The computation of the ideal 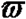 is illustrated (Fig. 3A). The example on the left indicates a situation where variability of the prior distribution is relatively smaller than the variability of the likelihood function. In this case, the ideal subject would “trust” the prior more by assigning smaller 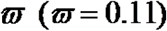. By contrast, the example on the right illustrates a situation where subjects should weigh likelihood information more heavily than prior 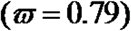 because variability of the likelihood function is relatively smaller than the variability of the prior distribution. It should be noted that the ideal 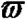 changes as a function of both the variability of the prior distribution and the sample size of the likelihood information (Fig. 3B).

**Figure 3.**
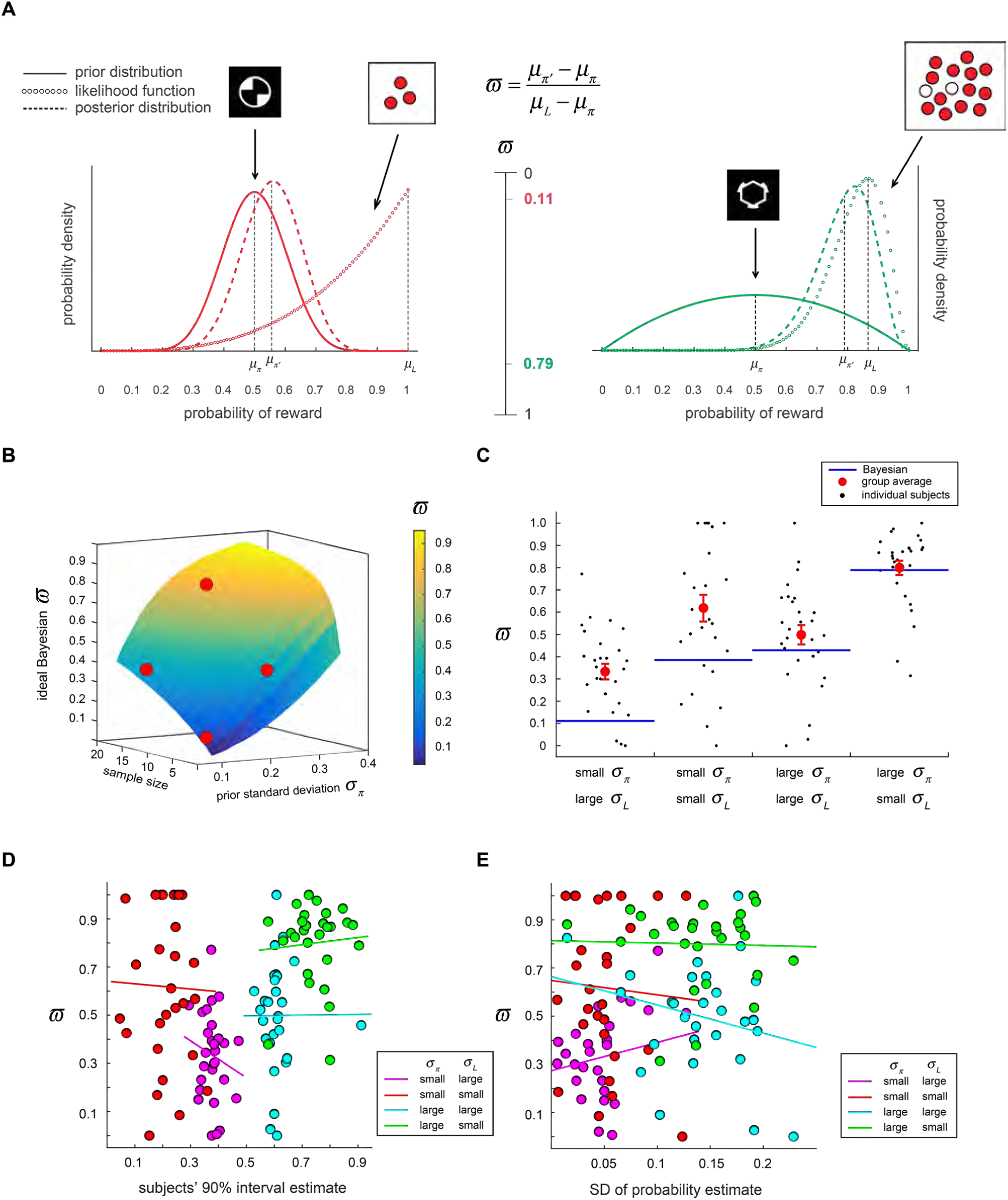
Behavioral results: Prior-likelihood integration session (Session 2). **A**. Two examples illustrating relative-weight computation 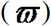 of the ideal Bayesian decision maker based on prior and likelihood information about probability of reward. **B**. Landscape of the ideal 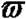 is plotted as a function of the variability of prior information (prior standard deviation) and variability of likelihood information (sample size). The four red dots indicate the combinations of prior and likelihood variability used in this experiment and their corresponding ideal 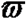. **C**.Subjective 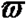 estimated based on subjects’ choice behavior compared with ideal 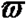 (blue lines). Data points in black indicate individual subjects’ 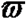 data points in red indicate mean of 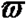 averaged across subjects. Error bars represent ±1 SEM. **D**. Subjective 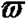 is plotted against standard deviation of subjects’ probability estimate from prior-learning session (Session 1). **E**. Subjective 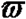 is plotted against subjects’ estimated 90 % interval from prior-learning session.

We found that subjects clearly adjusted 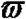 in response to likelihood variability – as likelihood variability increased 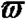 decreased (Fig. 3C). Subjects also adjusted 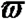 in response to prior variability – as prior variability increased 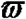 also increased. These results were qualitatively consistent with the direction predicted by the ideal Bayesian solution. Notably, we found that, compared with likelihood variability, subjects showed smaller adjustment in 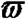 in response to changes in prior variability (t=−2.14, df=27, p <0.05).

When we compared subjective 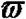 with the ideal Bayesian 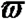 (see Behavioral analysis 2: ideal decision maker analysis in Methods), we found both near-optimal and suboptimal performance. When prior variability was large, mean subjective 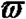 did not differ from the ideal 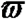 regardless of likelihood variability. By contrast, when prior variability was small, mean subjective 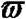 was significantly larger than ideal 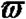, indicating that subjects underweight prior information. This pattern was seen regardless of likelihood variability. Further, this suboptimal behavior – underweighting of prior – cannot be attributed to individual differences in the variability estimate of the prior distributions in the prior-learning session: subjective 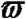 was not correlated with the standard deviation of subjects’ probability estimate (Fig. 3D); 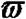 was also not correlated with the 90 % interval estimate (Fig. 3E) subjects provided in the prior-learning session. Together with the result showing that subjects had accurate knowledge about prior variability (Fig. 2B), these results indicated that weighting the prior and knowledge about the prior are dissociable – one could have very accurate knowledge about prior but still exhibits suboptimal weighting.

In summary, subjects did change 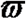 in response to changes in prior and likelihood variability in the direction consistent with Bayesian integration. However, subjects showed robust suboptimal integration by underweighting prior information. Such underweighting resulted from insufficient changes in weighting prior information in response to changes in prior variability but not to changes in likelihood variability. This phenomenon is a form of base-rate neglect widely reported in the literature (Kahneman & Tversky, 1973), but the degree of neglect was smaller compared with some classical results. Unlike a total neglect of prior information reported in Tversky and Kahneman (1974), subjects in our experiment did not show a complete neglect of prior information, which is consistent with Grether (1980).

### Orbitofrontal cortex and medial prefrontal cortex represent subjective weight of information

We used two separate fMRI analyses to identify neural representations of subjective 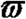. First, in a General Linear Model (GLM-1 in Methods) where trial-by-trial subjective 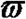 was implemented as a parametric regressor, we found significant activity in OFC and mPFC correlated with it (Fig. 4A, Table 1). Second, in a different GLM (GLM-2 in Methods) where we separately estimated average activity of each condition (a combination of prior and likelihood variability), we also found these two regions to correlate with subjective 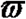 (whole-brain results in Fig. 4B). The central OFC (cOFC) correlated with the mean of subjective 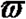 across four different conditions (ROI analysis in Fig. 4C), suggesting that cOFC represents changes in subjective 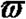 in response to changes in prior and likelihood variability. In addition, both cOFC and mPFC correlated with individual subjects’ 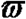 (Fig. 4DE). This indicates that for subjects who assigned larger 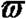 his or her activity in these regions tended to be larger. Together, these results indicate that while both cOFC and mPFC represent individual differences, but only cOFC tracked both the individual difference and the mean changes in 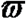 between different conditions.

**Table 1.**
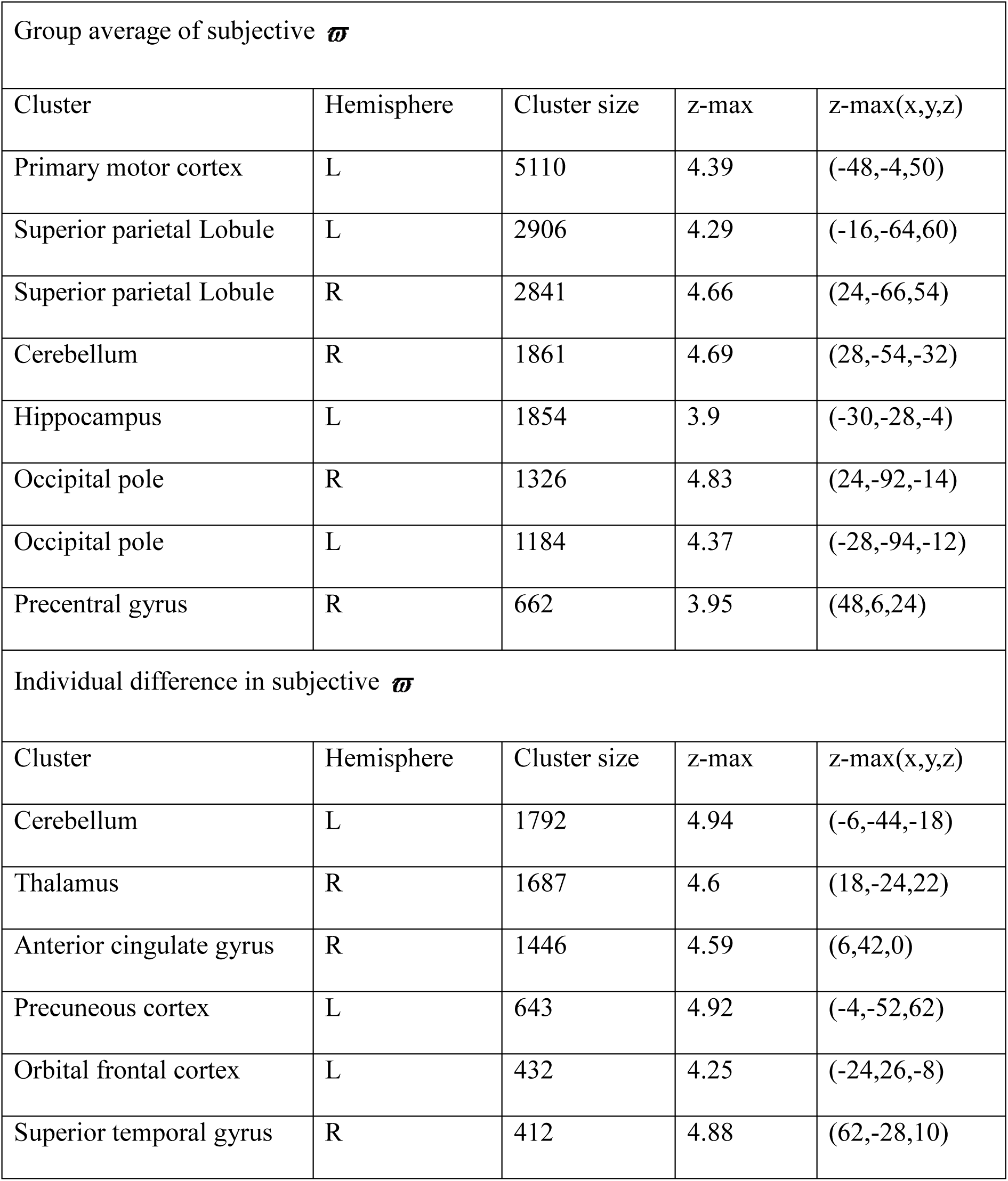
Regions in which the BOLD signal was positively correlated with the group average of subjective 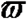 and the individual difference in subjective 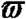. Clusters of Group average subjective 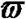 that survived cluster-based correction (z >2.4, family-wise error corrected at p <0.05 using Gaussian random field theory) and clusters of individual difference in subjective 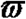 that survived cluster-based correction (z >3.1, family-wise error corrected at p <0.05 using Gaussian random field theory). The z-max column represents the MNI coordinates of the maximum z-statistic.

| Group average of subjective $\varpi$ | | | | |
| --- | --- | --- | --- | --- |
| Cluster | Hemisphere | Cluster size | z-max | z-max(x,y,z) |
| Primary motor cortex | L | 5110 | 4.39 | (-48,-4,50) |
| Superior parietal Lobule | L | 2906 | 4.29 | (-16,-64,60) |
| Superior parietal Lobule | R | 2841 | 4.66 | (24,-66,54) |
| Cerebellum | R | 1861 | 4.69 | (28,-54,-32) |
| Hippocampus | L | 1854 | 3.9 | (-30,-28,-4) |
| Occipital pole | R | 1326 | 4.83 | (24,-92,-14) |
| Occipital pole | L | 1184 | 4.37 | (-28,-94,-12) |
| Precentral gyrus | R | 662 | 3.95 | (48,6,24) |
| Individual difference in subjective $\varpi$ | | | | |
| Cluster | Hemisphere | Cluster size | z-max | z-max(x,y,z) |
| Cerebellum | L | 1792 | 4.94 | (-6,-44,-18) |
| Thalamus | R | 1687 | 4.6 | (18,-24,22) |
| Anterior cingulate gyrus | R | 1446 | 4.59 | (6,42,0) |
| Precuneous cortex | L | 643 | 4.92 | (-4,-52,62) |
| Orbital frontal cortex | L | 432 | 4.25 | (-24,26,-8) |
| Superior temporal gyrus | R | 412 | 4.88 | (62,-28,10) |

**Figure 4.**
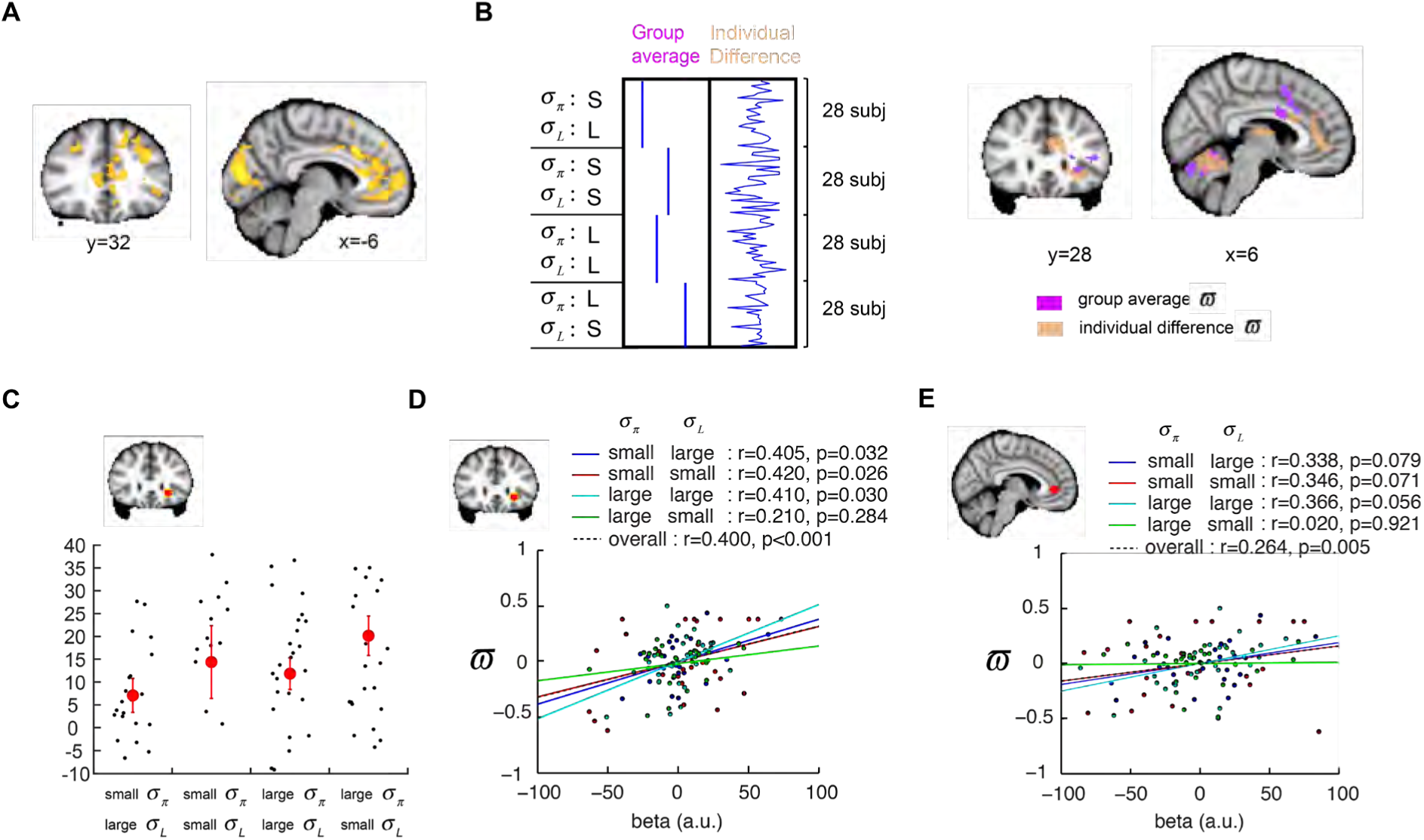
Orbitofrontal cortex (OFC) and medial prefrontal cortex (mPFC) represent subjective weight of information at the time of prior-likelihood integration. **A**. OFC and mPFC represent trial-by-trial subjective weight 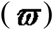 of information. **B**. Illustration of group-level covariate analysis that separately implements average subjective 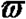 and individual differences in subjective 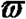 as parametric regressions. Whole-brain results showing regions that significantly correlated with mean 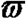 (magenta) and individual difference in 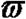 (light brown). **C**. Independent ROI analysis on cOFC. Mean activity (beta; average across subjects) of each condition is plotted (data points in red). The activity pattern closely resembles behavioral result shown in Fig. 3C. **D-E**. OFC and mPFC ROIs represent individuals’ subjective 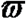. We plot subjective 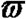 against brain activity in cOFC (4D) and mPFC (4E). Each data point represents a single subject’s data in a particular condition. Data points from different conditions are coded by different colors; the correlation between brain activity and subjective 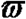 is shown for each condition separately. Overall correlation indicates correlation computed using all data points. Error bars represent ±1 SEM.

### Dorsal anterior cingulate cortex represents variability of prior and likelihood information

In a GLM (GLM-3 in Methods) where variability of prior and likelihood information were trial-by-trial parametric regressors, we found many brain regions that correlate, both positively and negatively, with information about prior and likelihood variability (Table 2). It is challenging to interpret these findings because any significant correlation, whether positive or negative, carries potentially important information. To tackle this problem, we used subjects’ behavioral performance, i.e. subjective 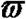 as a constraint for interpretation such that only the correlations that are relevant to subjective 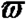 are considered meaningful. We know how 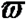 changed in response to changes in prior and likelihood variability (Fig. 3C): 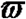 became larger when prior variability increased and when likelihood variability decreased (sample size from 3 dots to 15 dots). Prior variability therefore positively correlates with 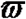, while likelihood variability negatively correlates with 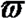 (Fig. 5A). Because of these relationships, we focused on regions that positively correlate with prior variability and negatively correlate with likelihood variability.

**Table 2.** Regions in which the BOLD signal was positively correlated with the standard deviation (SD) of the prior distribution (prior variability) and the SD of the likelihood information (likelihood variability). Clusters that survived cluster-based correction (z >2.5, family-wise error corrected at p <0.05 using Gaussian random field theory). The z-max column represents the MNI coordinates of the maximum z-statistic.

| Positive correlation with prior SD |  |  |  |  |
| --- | --- | --- | --- | --- |
| Cluster | Hemisphere | Cluster size | z-max | z-max(x,y,z) |
| Occipital cortex | L | 2344 | 3.69 | (-10,-80,0) in V1v |
| Positive correlation with likelihood SD |  |  |  |  |
| Cluster | Hemisphere | Cluster size | z-max | z-max(x,y,z) |
| Lateral Occipital cortex | L | 10075 | 5.02 | (-30,-76,16) |
| Lateral Occipital cortex | R | 2436 | 5.25 | (26,-66,54) |
| Prefrontal cortex | R | 1549 | 4.42 | (26,46,20) |
|  | R | 575 | 3.44 | (14,-24,42) |
| mPFC local maximum |  |  |  |  |
|  | R |  | 4.59 | (6,42,0) |
|  | L |  | 4.48 | (-4,30,24) |
|  | R |  | 4.45 | (6,40,6) |
|  | L |  | 4.31 | (-12,26,20) |
|  | R |  | 4.3 | (22,36,24) |
|  | L |  | 4.3 | (-10,28,26) |

| Negative correlation with likelihood SD |  |  |  |  |
| --- | --- | --- | --- | --- |
| Middle frontal Gyrus | R | 4870 | 4.99 | (46,18,48) |
| Lateral Occipital cortex | R | 1609 | 5.6 | (48,-56,42) |
| Lateral Occipital cortex | L | 1329 | 4.89 | (-50,-58,40) |
| Frontal Pole | L | 1002 | 4.36 | (-38,48,-14) |
| Frontal Pole | R | 780 | 5.18 | (36,44,-16) |
| Inferior temporal | R | 773 | 4.06 | (64,-38,-8) |
| Posterior Cingulate | L | 527 | 4.37 | (0,-30,34) |
| Precuneous Cortex | L | 476 | 4.06 | (-6,-74,38) |

**Figure 5.**
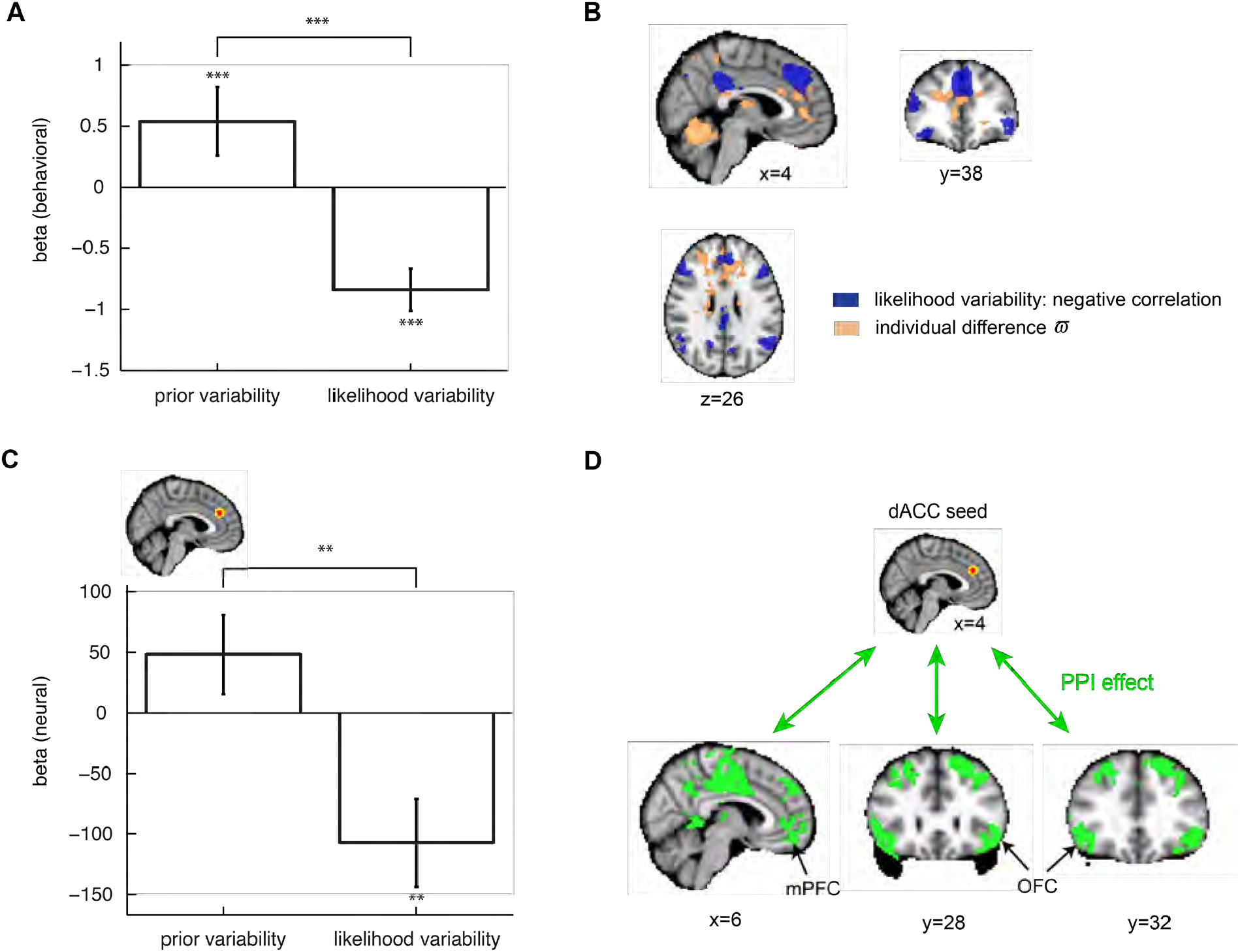
dACC represents variability of prior and likelihood information. **A**. Behavioral regression results showing that subjective 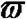 positively correlated with prior variability but negatively correlated with likelihood variability. **B**. Whole-brain results showing that part of dACC represents individual differences in subjective weight (blue) and negatively correlates with likelihood variability (light brown). **C**. Independent region-of-interest (ROI) analysis showed that dACC positively correlates with prior variability and negatively correlates with likelihood variability. **D**. Increased functional connectivity between variability-coding region in dACC and subjective-weight regions at the time of prior-likelihood integration. In a psychophysiologic interaction (PPI) analysis with dACC as seed region, we found a large number of regions (green), including OFC and mPFC that represented subjective 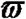, showed significant increase in functional connectivity with dACC at the time of prior-likelihood integration. * represents p <0.05; * * represents p <0.01; * * * represents p <0.001.

At the whole-brain level, there were regions that significantly and negatively correlated with likelihood variability (Fig. 5B in purple), but no region showed significant positive correlation with prior variability. Interestingly, we found that a region in the dorsal anterior cingulate cortex (dACC) that negatively correlated with likelihood variability partially overlapped with regions in mPFC that represented individual differences in 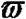. This prompted us to perform an ROI analysis: we constructed the ROI in dACC based on conjunction of significant negative correlation with likelihood variability (Fig. 5B in purple) and significant positive correlation with individual differences in 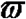 (Fig. 5B in light brown). For each subject, we created a sphere ROI using a leave-one-subject-out approach (see Methods for details), which led to slightly different spatial locations across different subjects. To visualize this, the voxels are coded in either red (overlap across all subjects) or yellow (otherwise) (Fig. 5C). The ROI results partially satisfied our criteria: it positively correlated with prior variability (marginally different from 0, p=0.1576) and negatively correlated with likelihood variability (p <0.01) (Fig. 5C, ROI results). Having said that, the difference in beta estimates between prior and likelihood variability was significant (p <0.01).

### dACC, OFC and mPFC constitute a functional network for prior-likelihood integration on reward probability

Based on the findings described above, we hypothesize that dACC, mPFC and OFC form an interconnected network that computes subjective 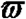 by using information about prior and likelihood variability. To test this hypothesis, we performed a psycho-physiologic interaction (PPI) analysis (Friston et al., 1997) using dACC as the seed region. We found that at the time of prior-likelihood integration, there was indeed an increase in functional connectivity between dACC and mPFC and OFC (Fig. 5D, Table 3). Although there are many other regions that also show increased connectivity with dACC, including the posterior cingulate cortex and the dorsolateral prefrontal cortex, these regions were not involved in representing subjective 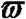.

**Table 3.**
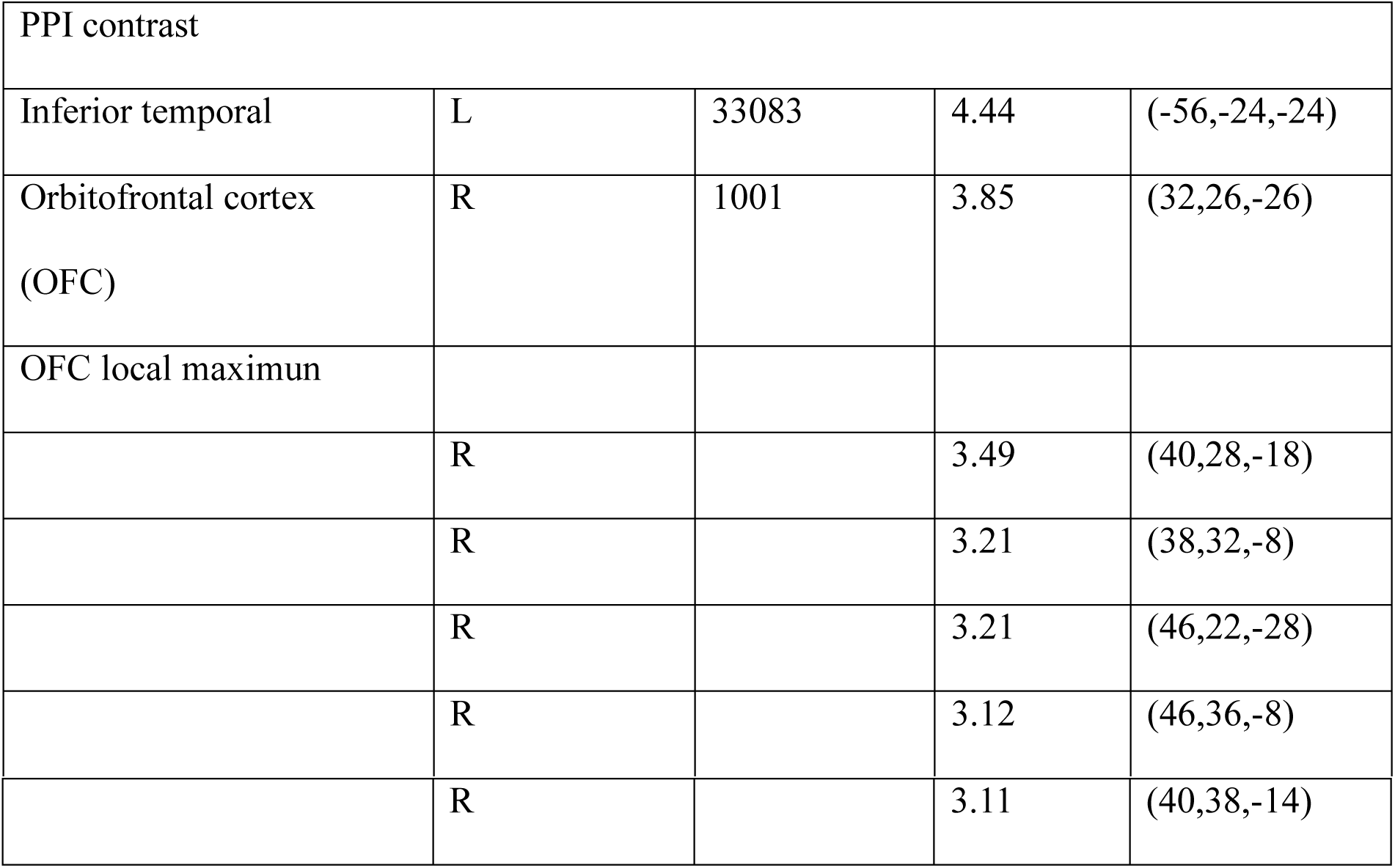
PPI results (use dACC as a seed) Clusters that survived cluster-based correction (z >2.3, family-wise error corrected at p <0.05 using Gaussian random field theory). The z-max column represents the MNI coordinates of the maximum z-statistic.

| PPI contrast |  |  |  |  |
| --- | --- | --- | --- | --- |
| Inferior temporal | L | 33083 | 4.44 | (-56,-24,-24) |
| Orbitofrontal cortex<br>(OFC) | R | 1001 | 3.85 | (32,26,-26) |
| OFC local maximum |  |  |  |  |
|  | R |  | 3.49 | (40,28,-18) |
|  | R |  | 3.21 | (38,32,-8) |
|  | R |  | 3.21 | (46,22,-28) |
|  | R |  | 3.12 | (46,36,-8) |
|  | R |  | 3.11 | (40,38,-14) |

### mPFC represents subjective value at the time of choice

In the GLM (GLM-1 in Methods) that included subjective value of the chosen option at the time of choice as parametric regressor, we found that mPFC significantly correlate with it (Fig. 6). Critically, the subjective value of the symbol lottery was the weighted sum of prior and likelihood information (Eq. 4 in Methods) where the weights directly came from subjective 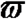. This not only replicated many previous studies showing that mPFC represents subjective value in value-based decision making, but also revealed that when multiple sources of information, such as prior and likelihood here, are present, mPFC first computes relative subjective weight assigned to these sources of information and then uses it to compute subjective value. In summary, we found that mPFC is critically involved in both probabilistic inference and decision making (Fig. 6): at the inference stage, mPFC activity represents subjective weight of information critical to the computation of subjective value. At the time of choice, mPFC represents the subjective value of the chosen option.

**Figure 6.**
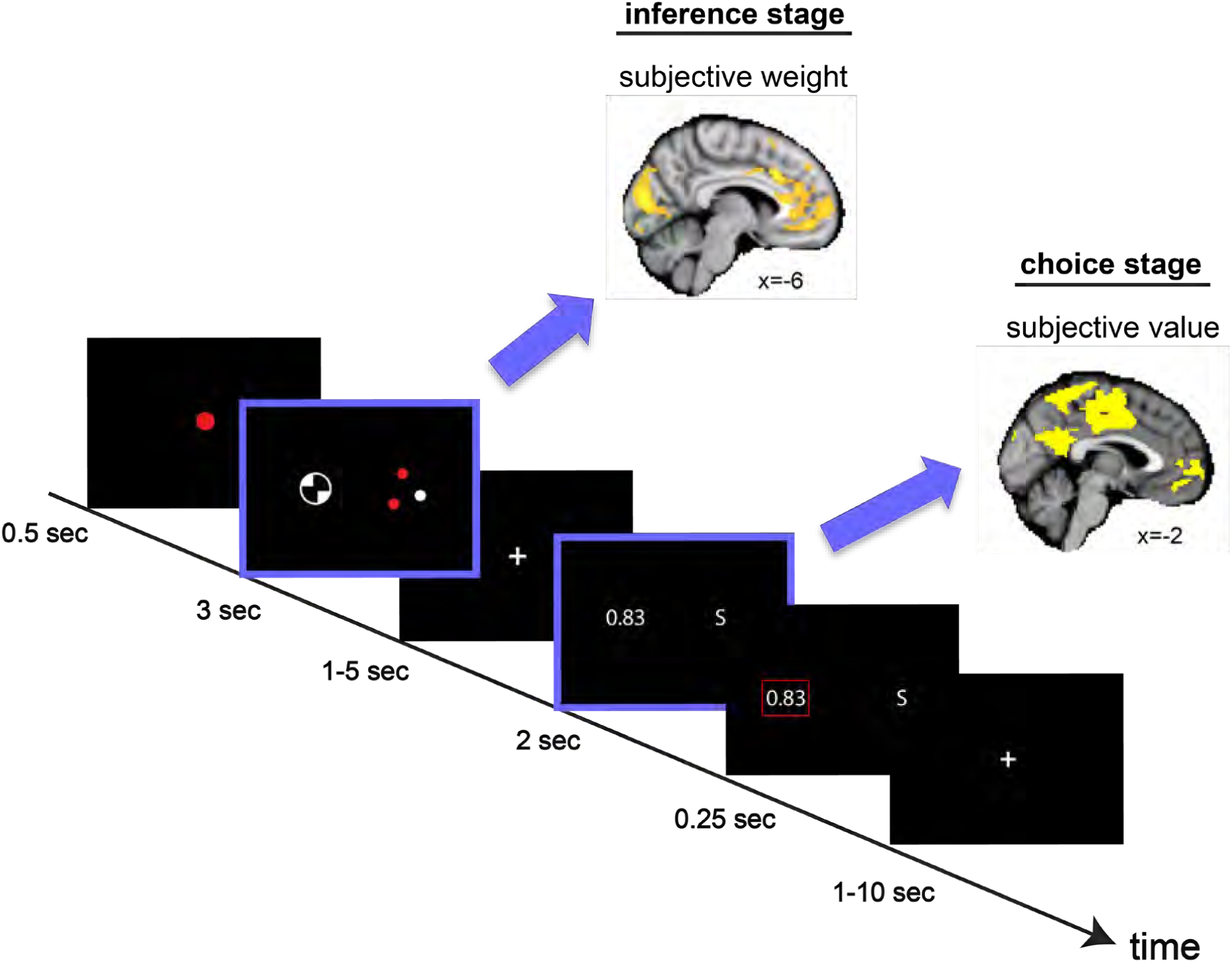
Medial prefrontal cortex (mPFC) is involved in probabilistic inference and decision making. Here we show that, at the time of inference, mPFC represents subjective 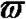 critical to the integration of prior and likelihood information. At the time of choice, mPFC represents subjective value of the chosen option.

## Discussion

Uncertainty is a central feature in many decisions we face. To make decisions under uncertainty, we must estimate probability of uncertain events based on available information. Previous research showed that humans exhibit systematic biases in probability estimation. In this study, we investigated the neuraocomputational basis of a well-known bias, the base-rate neglect, in which people underweight base rate or prior information (Kahneman, Slovic, & Tversky, 1982). We found robust base-rate neglect despite strong evidence that subjects indeed combined prior and likelihood information in the direction consistent with an ideal Bayesian integrator. By independently manipulating variability of prior and likelihood information, we were able to establish that base-rate neglect largely resulted from insufficient changes in weighting prior information in response to changes in the variability of prior information, but not to changes in the variability of the likelihood information. Interestingly, independent of likelihood variability, subjects showed base-rate neglect when they should “trust” prior more – when prior variability was small. Together, these results provide insights into base-rate neglect by identifying prior variability as the critical statistical attribute contributing to this bias.

### Neural computations for subjective weight

In our study, we found that OFC and mPFC represent the subjective weight subjects assigned to prior and likelihood information about probability of reward. mPFC and OFC not only correlated with individual differences in subjective weight but also showed trial-by-trial representation of subjective weight. Vilares et al. (2012) reported similar findings (trial-by-trial subjective weight termed Bayesian instantaneous slope) in the superior medial prefrontal cortex in a visuo-motor inference task. Together, these results indicate that mPFC is crucial in computing subjective weight.

The finding on subjective weight connects closely with the value-based decision literature, which show that OFC and mPFC are part of a valuation network that computes subjective value of goods (Kable & Glimcher, 2009; Padoa-Schioppa, 2011). Subjective value is an integrated signal that summarizes the desirability of an option through combining different attributes, variables or sources of information associated with an option. In an inference or estimation task such as the one used in this or in multiattribute choice, subjective value is often modeled as the weighted sum of different sources of information or attributes where subjective weights specify the weights a decision maker assigns to these different sources or attributes. In this study, subjective weights are necessary to estimating probability of reward, which is computed as the weighted sum of prior probability of reward (the mean of the prior distribution) and likelihood of reward. In other words, without subjective weights, it would not be possible to determine how prior and likelihood information are combined to estimate probability of reward. Our finding therefore established the evidence that subjective weight – precursor to computing subjective value – is represented in OFC and mPFC. This is consistent with the findings that OFC represents different features or attributes of value information (Blanchard, Hayden, & Bromberg-Martin, 2015; O’Neill & Schultz, 2010) and represents the subjective weight of attributes in multiattribute choice (Hare, Camerer, & Rangel, 2009).

Growing evidence indicates that mPFC is heavily involved in inference through combining different sources of information, in particular, the integration of prior and likelihood. In an intention inference task where subjects were asked to infer the intention of a person’s action, Chambon et al. (2017) found that mPFC correlated with the interaction of subjects’ prior belief on intention and the strength of sensory evidence. The interaction indicates that mPFC activity represents the integration of prior and likelihood information about intention. In Chan et al. (2016), subjects were asked to infer latent causes in an inference task where probability distribution on latent causes were manipulated. They found that multivoxel pattern of activity in OFC represents the log posterior probability distribution on latent causes. In our previous study (Ting et al., 2015), we found that mPFC correlated with the mean of the posterior distribution on probability of reward. The present study adds to this literature by showing that mPFC represents the subjective weight necessary for the integration of prior and likelihood information when these two sources of information were presented. We also found that mPFC represents the subjective posterior probability of reward – the weighted sum of prior and likelihood information that takes into account subjective weight. However, the subjective posterior appeared at a later stage of the trial – presumably after subjective-weight computations – where subjects had use subjective posterior to make a choice between different options.

### Temporally dissociable representations of subjective weight and subjective value

In this study, we found that subjective-weight and subjective-value representations in OFC and mPFC are temporally dissociable. By temporally separating inference (where subjects combined prior and likelihood information to estimate probability of reward) from choice (where subjects had to use his probability estimate to make a choice) through task design, we found that during the inference stage activity in these two regions represent subjective weight. By contrast, at the time of choice, mPFC and OFC represents subjective value of the chosen option. That is, when the subjects chose the symbol lottery, subjective value was its subjective posterior reward probability; when alternative lottery was chosen, subjective value was its probability of reward. We used probability of reward to represent subjective value because reward magnitude was fixed throughout the experiment. This finding indicates that mPFC and OFC represent subjective weight and subjective value at two different points in time. Further, the order of representation (subjective weight first followed by subjective value) suggests that these two regions first compute the weights that the decision maker later assigns to prior and likelihood when computing the subjective value of the option.

### Dissociable representations of information variability and subjective weight

In contrast to the OFC and mPFC findings on subjective weights, we found that dACC represents variability of prior and likelihood information. Previous studies showed that dACC represents variability or degree of uncertainty in receiving a reward (Critchley, Mathias, & Dolan, 2001) and variance in the reward distribution (Burke & Tobler, 2011; Christopoulos, Tobler, Bossaerts, Dolan, & Schultz, 2009). In our study, variability of prior and likelihood information indicate variability in the probability of reward, i.e. the standard deviation of the prior distribution and likelihood function on probability of reward. Together with previous findings, these results seem to indicate a general role of dACC in representing variability of information that can be formally described by the variance statistic of a distribution.

Further, our study showed that the direction of variability coding in dACC carries behavioral significance. We found that dACC positively correlated with prior variability and negatively correlated with likelihood variability. The signs of correlation match with how subjective weight correlated with prior and likelihood variability. Because of such psychometric-neurometric match, we interpret variability-coding in dACC to be tightly linked with subjective-weight computations in mPFC and OFC. This is further supported by the finding that at the time of prior-likelihood integration, dACC exhibited an increase in functional connectivity with mPFC and OFC. This suggests that variability-coding in dACC interacts with mPFC and OFC potentially for the purpose of determining subjective weight based on variability of prior and likelihood information.

In summary, the present study provided crucial evidence for the neurocomputational substrates and neural mechanistic account for base-rate neglect. We showed that base-rate neglect arises from insufficient adjustment in weighting prior information in response to changes in the variability of prior information. The neurocomputational substrates of base-rate neglect appear to be in OFC and mPFC, as these two regions represent subjective weight that reflects base-rate neglect. Subjective-weight representations in OFC and mPFC further interact with variability-coding region in dACC at the time of prior-likelihood integration, suggesting that the interactive and connecting properties of these regions play an important role in base-rate neglect. In the future, it would be important to identify mechanisms that could change the degree of base-rate neglect, i.e. ways that would change the degree of adjustment in weighting prior information and to characterize how potential behavioral changes affect the representations of subjective weight and the interactions between subjective-weight network (OFC and mPFC) and variability-coding regions such as dACC.

## Materials and Methods

The data and analysis code are available at https://osf.io/ku97p/.

### Subjects

Twenty-eight subjects participated in the experiment (fourteen males; mean age = 24.6 years; age range = 21-30 years) and completed two sessions on two consecutive days. All participants had no psychiatric or neurological disorders. All subjects gave written informed consent prior to participation in accordance with the procedures approved by Taipei Veterans General Hospital IRB. Subjects were paid 620 New Taiwan Dollar (NTD, 1 US Dollar=30 NTD) for their participation and additional monetary bonus (average: 383 NTD) based on their performance in the experiment.

### Procedure

We designed an experiment to investigate how humans combine prior knowledge and current (likelihood) information about probability of reward and the neural systems involved in performing such integration computation. There were two sessions in the experiment. The first session was conducted in a behavior testing room. The second session – the primary focus of the experiment – was conducted in an MRI scanner. The tasks were programmed using the Psychophysics Toolbox in MATLAB (Brainard, 1997; Pelli, 1997).

### Session 1 (behavior): learning prior distributions

The goal of this session was to establish knowledge about probability of reward associated with different visual stimuli. There were two visual stimuli, each representing a unique probability density function on probability of reward. Both were beta distributions with two parameters *α* and *β*. Critically, we manipulated the variability of prior knowledge by varying the variance of density functions. The standard deviation of the two prior distributions was 0.1, (*α,β*)=(12,12), and 0.2236, (*α,β*)=(2,2). We use *σ*_*π*_ to denote the standard deviation of the prior distribution. The mean of these two prior distributions were kept at 0.5. This indicated that both stimuli had an average of 50 % chance of receiving a monetary reward. The magnitude of reward was fixed throughout the experiment.

Subjects did not know the probability distributions associated with the two stimuli before the experiment and had to acquire this knowledge through sampling from them. There were 10 blocks of trials, each consisting of 30 trials. Within each block, only one of the two visual stimuli was presented. The small-*σ*_*π*_ stimulus was presented in five blocks and the large-*σ*_*π*_ stimulus was presented in the remaining 5 blocks. The ordering of the blocks was semi-randomized so that subjects encountered no more than 2 successive blocks with the same variance.

On each trial, the subjects were instructed to estimate probability of reward associated with the presented visual stimulus, which was sampled from its corresponding probability density function. Subjects indicated his/her probability estimate by pressing number keys on a computer keyboard. For example, subjects would enter 7 followed by 2 and then press the Enter key if she or he believed that the probability of reward was 72 %. After indicating the probability estimate, the true probability of reward on that trial was revealed. Note that since the probability of reward on each trial was drawn from a probability density function, the probability of reward would vary from trial to trial. How much it varied would depend on the variance of the distribution.

In order to motivate the subjects to learn the probability distribution, we implemented an incentive compatible procedure as in Ting et al. (2015). On each trial, subjects received a monetary gain or loss depending on how close his/her estimate on the probability of reward was to the true probability of reward. Subjects would receive 100 points if the difference between estimated and true reward probability was within 5 %, 50 points if the difference was between 6 to 10 %, 0 if the difference was between 10 % and 20 %, and would lose 50 points if the difference was greater than 20 % (100 points = 1 NTD). Moreover, to examine whether subjects acquired knowledge about the mean and variance of the two prior distributions, at the end of each block, subjects were asked to estimate the mean and the 90 % confidence interval of the probability of reward associated with the stimulus s/he encountered in that block.

### Session 2 (fMRI): integrating prior and likelihood

The session was conducted on the day after Session 1. Its goal was to investigate the neural mechanisms for integrating prior knowledge – established through Session 1 – and the likelihood information about probability of reward. Below we describe a lottery decision task designed to address this question.

On each trial, the subjects were asked to choose between two lotteries that differed only in the probability of receiving a small monetary reward. The magnitude of reward associated with both options was the same and fixed throughout the experiment so that subjects should make their decision based on which option carried a larger probability of reward. For one of the lottery options, referred to as the symbol lottery, its probability of reward was not explicitly stated in numeric or graphical forms – subjects needed to infer the probability based on two pieces of information, prior and likelihood. On each trial, prior information was represented by one of the two visual stimuli (a symbol icon) that the subjects encountered in Session 1. Each stimulus represented a probability density function on probability of reward. Subjects were told the probability of reward associated with the symbol lottery was sampled from the density function. Meanwhile, the likelihood information was represented by a set of colored dots (red or white). Given the probability of reward associated with the symbol lottery on the current trial, the dots summarized the sample drawn from it. Each red dot represented a reward outcome and each white dot represented a no-reward outcome. Hence, the proportion of red dots indicates the likelihood of reward. Note that if the sample size were infinitely large, the proportion of red dots would be equal to the probability of reward.

We manipulated the variability of the likelihood information by varying the sample size (number of dots: 3 or 15) used to draw from the probability of reward. Together with the manipulation of the variance of prior distribution, we achieved a 2 (prior variability, *σ*_*π*_, small and large) × 2 (likelihood variability, *σ*_*L*_, small and large) ×2factorial design. We use *σ*_*π*_ and *σ*_*L*_to denote the standard deviation of prior and likelihood function respectively. We refer to each combination of prior and likelihood variability as a condition. The average standard deviation of the likelihood function for the smaller sample size (3 dots) and larger sample size (15 dots) was 0.2722 and 0.1205 respectively. These values are almost identical to the standard deviation of the prior information (0.1 and 0.2236). As a result, the difference in the range of variance between the prior and likelihood information was controlled.

On each trial, the prior and likelihood information were presented on the left and right side of the screen, with the locations randomized across trials. Following the presentation of the symbol lottery, there was a fixation period (1 ∼5 seconds, discrete uniform distribution in steps of 1 second). This was followed by the presentation of the second lottery, also referred to as the alternative lottery. Information about its probability of reward was explicitly revealed in number form. In Fig. 1C, as an example, the alternative lottery has a 0.83 probability of reward. The reward probability of the alternative lottery was determined in the following way. In order to reliably estimate the weight the subjects assign to prior and likelihood information about the symbol lottery, we designed the reward probability of the alternative lottery such that on each trial it was drawn randomly from a uniform distribution (*μ*_*π’*_ –0.4, *μ*_*π’*_ +0.4) where *μ*_*π’*_ denotes the mean of ideal posterior distribution associated with the symbol lottery on the current trial (see Bayesian integration model below for details). With this design, first, we expected the choices that subjects faced to be non-trivial – the alternative lottery on average had a probability of reward that should be close to the subjects’ estimated probability of reward for the symbol lottery if subjects were ideal. Even if subjects were not ideal but nonetheless performed prior-likelihood integration in a direction consistent with the ideal integration, his or her probability estimate should be within this range. Second, we expected that subjects would not be able to predict the probability of reward associated with the alternative lottery at the time of prior-likelihood integration (when information about prior and likelihood were revealed). This had several advantages: (1) it would be harder for the subjects to derive a fixed decision strategy; (2) it kept subjects engaged in the decision task, which should motivate them to combine prior and likelihood information throughout the experiment; (3) for analysis of fMRI data that focused on the period where prior-likelihood integration took place (when information about prior and likelihood were revealed), not being able to know the alternative lottery prevented the subjects from comparing the two lotteries at the time of prior-likelihood integration. This allowed us to dissociate prior-likelihood integration from possible confounds such as value comparison or motor preparation in the fMRI analysis that focused on the time when information about prior and likelihood were revealed. Finally, to avoid the possibility of getting extremely small or extremely large probabilities of reward for the alternative lottery, we further constrained the probability of reward to be between 0.01 and 0.99. That is, on each trial, we randomly sampled within the range of max[*μ*_*π’*_ –0.4,0.01]and max[*μ*_*π’*_ +0.4,0.99]to determine the probability of reward associated with the alternative lottery.

When the alternative lottery was presented, subjects were instructed to choose between the symbol lottery (indicated by S) and the alternative lottery (a number) within 2 seconds. The location of the lotteries (left or right) was randomized and balanced across trials. Once the subjects indicated his or her decision with a button press, the chosen option was revealed on the screen for 250ms. No feedback on the outcome of the chosen option was revealed. This was to prevent the subjects from updating knowledge about the prior distribution associated with the visual stimulus and learning how to integrate prior and likelihood information through feedback.

There were 6 blocks in the session. Each block had 40 trials. Each combination of the prior symbol icon (high prior variability, low prior variability) and sample size (3 dots, 15 dots) for the likelihood information had 10 trials in each block. The order of the trials was randomized.

#### Mathematical notation

We summarize the notations used when describing behavioral and fMRI analyses below. We denote the mean of posterior distribution on reward probability associated with the symbol lottery as *μ*_*π’*_, the standard deviation of the posterior distribution as *σ*_*π’*_, the standard deviation of the prior distribution as *σ*_*π*_, the likelihood of reward (proportion of red dots) as *μ*_*L*_, the standard deviation of the likelihood function as *σ*_*L*_,, the probability of reward associated with the alternative lottery as *θ*_alt_, and the subjective weight the subjects assigned to the likelihood information relative to the prior information as 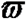.

### Bayesian integration model

Below we describe the ideal Bayesian integration of prior and likelihood information about the symbol lottery in the lottery decision task (Session 2). Let *θ* denote the probability of reward associated with the symbol lottery on a trial. The subjects did not know the true value of *θ* on each trial but could estimate it based on prior knowledge and likelihood information. Subjects were trained to learn two different prior distributions in a previous session (see Session 1: learning prior distributions). Both were beta distributions. Let *π*(*θ*) denote a prior distribution

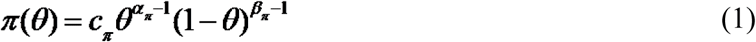

where ***c*** _*π*_ is a normalization constant, *α*_*π*_ and *β*_π_ are the free parameters. The likelihood function, *L*(*θ*) is also a beta distribution

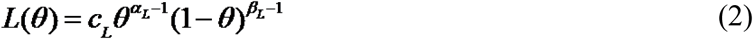

where *c*_*L*_ is a normalization constant, *α*_*L*_ and *β*_*L*_ are the free parameters. *α*_*L*_ and *β*_*L*_ have the following relation with the sample: let sbe the number of successes in the sample and *f* the number of failures. Then *α*_*L*_= *s* +1, *β*_*L*_ = *f* +1. Note that the sample – represented by colored dots on the screen – was drawn from the true value of *θ* on the particular trial. In the sample, winning a reward (a success) was represented by a red dot, while winning nothing (a failure) was represented by a white dot.

The posterior distribution, the product of the prior distribution and the likelihood function, is also a beta distribution (O’Hagan, Forster, & Kendall, 2004)

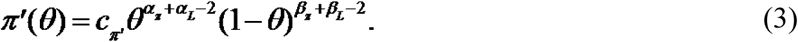

Let *α*_*π’*_ and *β*_*π’*_ represent the free parameters of the posterior distribution, then *α*_*π’*_= *α*_π_ + *α*_*L*_ − 1 and *β*_*π’*_= *β*_π_ + *β*_*L*_ –1 or equivalently *α*_*π’*_= *α*_π_ +s and *β*_*π’*_= *β*_π_ + *f*. The expected value (mean) of the posterior distribution (*μ*_*π*’_) is *α*_*π’*_/(*α*_*π’*_+*β*_*π’*_), which we take to be the estimate of *θ* of an ideal Bayesian integrator. An alternative is the mode – the maximum a posteriori (MAP) estimator, which in the beta distributions considered here, is very close to the mean and would lead to the same conclusion here. In this study, we used Eq.(3) to simulate choice data of an ideal Bayesian decision maker, estimated the ideal weight assigned to prior and likelihood information and compared it with the weight our subjects assigned to these two sources of information (see Behavioral analysis 3: ideal decision maker analysis below for details).

### Behavioral analysis 1: Estimating subjective weight

We developed a method for estimating subjective weight defined as the weight assigned to likelihood information relative to prior 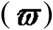 using subjects’ choice data and the ideal Bayesian integrator’s choice data based on simulations. In this paper, 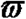 serves as the basis for comparing between an ideal Bayesian integrator and subjects’ integration performance and our goal was to investigate how and how well subjects adjusted subjective 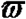 in response to changes in prior and likelihood variability. Therefore, for each subject and each condition (a combination of prior and likelihood variability) separately, we estimated subjective 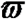 using choice data. We further compared subjective 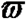 with the 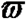 of an ideal Bayesian integrator facing exactly the same experiment as the subjects (see Behavioral analysis 2: ideal decision maker analysis).

We assumed that the subject’s estimate on the probability of reward associated with the symbol lottery 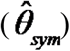 is a linear combination of *μ*_*π*_ and *μ*_*L*_

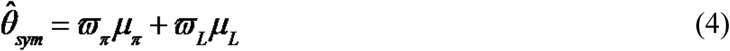

where 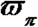 and 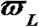 represent the weight assigned to *μ*_*π*_ and *μ*_*L*_ respectively. Since in this experiment *μ*_*π*_ was fixed (0.5 for both prior distributions used), it is effectively a constant. For this reason, we only estimated 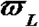 and assumed that 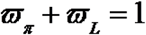 and that 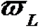 is between 0 and 1. Note that in this model, 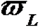 is relative in nature and should be interpreted as the weight assigned to the likelihood information relative to the prior information. For convenience, we further denote 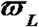 as 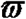 and rewrite Eq. (4)

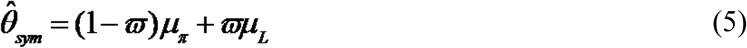

Given computed 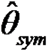 from Eq. (5), we then modeled the probability of choosing symbol lottery by

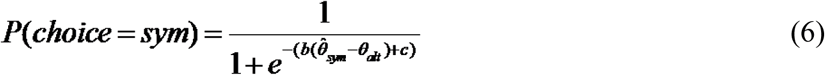

where *θ*_*alt*_ represents the probability of reward associated with the alternative lottery, b and c are the free parameters to be estimated. We used method of maximum likelihood to estimate three parameters – 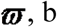, and c – for each subject and each condition separately.

Finally, note that even within the same condition, both *μ*_*L*_ (likelihood of reward represented by proportion of red dots) and *σ*_*L*_would vary from trial to trial and therefore 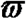 should also vary from trial to trial. However, separately estimating 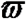 for each trial would make the parameter estimate extremely noisy, making this approach unrealistic. Given that sample size is the dominant source of variance of *σ*_*L*_, it is reasonable to estimate 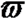 for each condition (instead of trial-by-trial estimate) separately. In this case, 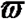 should be interpreted as the average 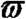 of across trials within a condition.

### Behavioral analysis 2: Ideal decision maker analysis

To understand whether the subjects optimally integrated prior and likelihood information, we simulated choice data of an ideal decision maker (using Eq. 3 to compute *μ*_*π*’_ and Eq. 5 to simulate choice) when facing exactly the same trials as the subjects faced. Our goal was to estimate the ideal 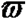 (Eq. 4) compared it with the subjects’ actual 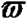. We simulated 10,000 experiments performed by an ideal decision maker and in each simulated experiment we estimated, for each of the 4 conditions separately, 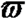 of the ideal decision maker (2 prior variability × 2 likelihood variability) using exactly the same procedure described in Behavioral analysis 1 described above. Finally, for each condition separately, we computed the mean of the ideal 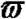 from 10,000 simulated experiments and used it as the ideal 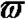. Interestingly, we also found that the ideal 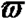 computed this way is identical to simply calculating 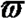 for each condition (a combination of prior variability and sample size) using the following equation

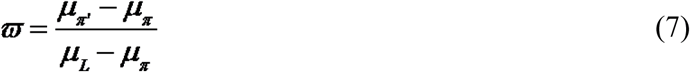

where *μ*_*π*_ is the mean of the prior distribution, *μ*_*L*_is the likelihood of reward and *μ*_*π*’_ is the mean of the posterior distribution. We also found that in a given condition, across different simulated trials where *μ*_*π*’_ and *μ*_*L*_ would vary, 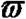 would always be the same. Finally, to compare the ideal 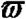 with the subjective 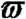 estimated from the subjects’ choice behavior, for each condition, we performed a t-test with the null hypothesis that the subjective 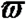 is equal to the ideal 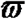.

#### fMRI data acquisition

Subjects completed the task in a 3T Siemens MRI scanner (MAGNETOM Trio) equipped with a 32-channel head array coil. Each subject completed six runs. Two localizer scans were implemented, one at the beginning of the experiment and the other midway through the experiment (after the first 3 runs). For each run, T2 *-weighted functional images were collected using an EPI sequence (TR=2000ms, TE=30ms, 33 oblique slices acquired in ascending interleaved order, 3.4 ×3.4 ×3.4 mm isotropic voxel, 64 ×64 matrix in 220 mm field of view, flip angle 90 °). To reduce signal loss in the ventromedial prefrontal cortex and orbitofrontal cortex, the sagittal axis was tilted clockwise up to 30 °. Each run consisted of 40 trials and 285 images. After functional scans, T1-weighted structural images were collected (TR=2530ms, TE=3.03ms, flip angle=7 °, 192 sagittal slices, 1 ×1 ×1 mm isotropic voxel, 224 ×256 matrix in a 256-mm field of view). For each subject, field map image was also acquired for the purpose of estimating and partially compensating for geometric distortion of the EPI image so as to improve registration with the T1-weighted images.

#### fMRI preprocessing

The imaging data were preprocessed with FMRIB’s software Library (FSL). First, for motion correction, MCFLIRT was used to remove the effect of head motion during each run. Second, FUGUE (FMRIB’s Utility for Geometric Unwarping of EPIs) was used to estimate and partially compensate for geometric distortion of the EPI images using field map images collected for the subject. Third, spatial smoothing was applied with a Gaussian kernel with FWHM=5mm. Fourth, a high-pass temporal filtering was applied using Gaussian-weighted least square straight line fitting with *σ*=50s. Fifth, registration was performed in a two-step procedure, with the field map used to improve the performance of registration. First, EPI images were registered to the high-resolution brain T1-weighted structural image (non-brain structures were removed via FSL’s BET (Brain Extraction Tool). Second, the transformation matrix (12-parameter affine transformation) from the T1-weighted image to the Montreal Neurological Institute (MNI) template brain was estimated. This allowed for transforming the EPI images to the standard MNI template brain.

### General Linear Models of BOLD signal

All GLM analyses were carried out in the following steps (Beckmann, Jenkinson, & Smith, 2003). First, BOLD time series were pre-whitened with local autocorrelation correction. A first-level FEAT analysis was carried out for each run of each subject. Second, a second-level (subject-level) fixed-effect (FE) analysis was carried out for each subject that combined the first-level FEAT results from different runs using the summary statistics approach. Finally, a third-level (group-level) mixed-effect (ME) analysis using FSL’s FLAME module (FMRIB’s Local Analysis of Mixed Effects) was carried out across subjects by taking the FE results from the previous level and treating subjects as a random effect (Woolrich, Behrens, Beckmann, Jenkinson, & Smith, 2004). All reported whole-brain results were corrected for multiple comparisons. We first identified clusters of activation by defining a cluster-forming threshold of the z statistic. Then, a family-wise error corrected p-value of each cluster based on its size was estimated using Gaussian random field theory (Worsley, Evans, Marrett, & Neelin, 1992).

#### GLM-1

In this model, we implemented the following regressors: on each trial, at the time of prior and likelihood presentation, (R1) an indicator function with a duration of 3s, (R2) R1 multiplied by subjective 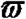, (R3) R1 multiplied by subjective posterior probability of reward associated with the symbol lottery (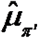) computed based on Eq. (5) given subjective weight 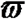, (R4) R1 multiplied by sample size (number of dots) of the likelihood information. This is to rule out the possibility that regions correlated with subjective-weight is due to sample size, which partially correlates with subjective weight. At the inter-stimulus interval (ISI, 1 ∼5sec, discrete uniform distribution in steps of 1 sec) between prior-likelihood presentation and choice, (R5) an indicator function with duration equal to the length of ISI of that trial. At the time of choice where alternative lottery was presented, (R6) an indicator function with duration of subject’s response time, (R7) R6 multiplied by the subjective value of the chosen lottery. For R7, when the chosen lottery was the symbol lottery, subjective value indicates the subjective posterior 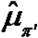; when the alternative lottery was chosen, subjective value was its probability of reward. Typically, subjective value is the integration of subjective reward magnitude with subjective probability. However, since reward magnitude was fixed throughout the experiment, subjective reward magnitude was effectively a constant and therefore we could use subjective probability to represent subjective value.

#### GLM-2

This model estimated mean activity in response to each of the four conditions (2 prior variability × 2 likelihood variability) at the time of prior-likelihood integration so that it can be used to correlate with subjective 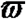 of each condition for each subject (see Group-level covariate analysis below). Hence, for each condition separately, at the time of prior and likelihood presentation, we implemented the following regressors: (R1) an indicator function with duration of 3s, (R2) R1 multiplied by *μ*_*L*_. At the time of choice (where alternative lottery was presented), we implemented (R9) an indicator function with duration of the subject’s response time, (R10) R9 multiplied by *θ*_*alt*_, (R11) R9 multiplied by *μ*_*π*’_, (R12) R9 multiplied by *σ*_*π’*_.

#### Group-level covariate analysis

We performed a group-level mixed-effect analysis to identify regions whose activity correlated with subjective 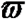. The inputs of the model were the beta estimate of the four indicator regressors (R1, one for each condition) in GLM-2 obtained at the second-level (subject-level fixed-effect model) analysis from each subject. For each condition, the beta estimate of the indicator regressor (R1 in GLM-2) reflects the mean BOLD response (over trials of the same condition) at the time when prior and likelihood information were presented. Because there were 28 subjects and four beta estimates per subject, there were a total of 112 (28 ×4) input beta images. Together, these inputs represented the estimated activity in response to the four different conditions at the time of information integration. We implemented the following regressors: (R1) an indicator function that modeled the mean activity (across all conditions), (R2) the mean of subjective 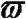 of each condition (averaged across all subjects) minus the mean of subjective 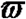 averaged across all subjects and conditions, (R3) subjective 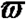 of each subject in a condition minus the mean of 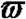 of that condition. We set up two contrasts – the contrast [R1 R2 R3]=[0 1 0] was used to identify the neural correlates of group mean of 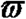 (mean 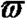 averaged across subjects) associated with the 4 different conditions, and contrast [R1 R2 R3]=[0 0 1] was used to examine the neural correlates of individual differences in 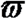 (the deviation of individuals’ 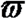 from the mean 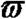 of each condition). The results of group average 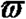 were corrected for multiple comparisons (z > 2.4, familywise error corrected at p <0.05 using Gaussian random field theory), and the results of individual 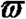 were corrected for multiple comparisons (z > 3.1, familywise error corrected at p <0.05 using Gaussian random field theory)

#### GLM-3

To identify regions that correlate with variability of prior and likelihood information at the time of their presentation, we implemented the following regressors: (R1) an indicator function with duration of 3s, (R2) R1 multiplied by *σ*_*π*_, (R3) R1 multiplied by *σ*_*L*_. In addition, we also implemented (R4) R1 multiplied by *μ*_*L*_ and (R5) R1 multiplied by sample size (number of dots) of the likelihood information. The regressors implemented at the time of choice were identical to the ones described in GLM-2. The analysis was corrected for multiple comparisons (z >2.5, family-wise error-corrected at p <0.05 using Gaussian random field theory)

#### Independent regions-of-interest (ROIs) analysis

Based on results from the whole-brain analysis, we created independent and unbiased ROIs using the leave-one-subject-out (LOSO) method (Litt, Plassmann, Shiv, & Rangel, 2011; Ting et al., 2015). The LOSO method was used to analyze *w*_*L*_ and representations for *σ*_*π*_ and *σ*_*L*_ described above. For each subject separately, we performed the analysis in the following steps. First, we identified regions that correlated with a contrast of interest (e.g. *σ*_*L*_) using all other subjects’ data. Second, we identified the voxel with the maximum z-statistic among the voxels that significantly correlated with the contrast of interest and created a sphere mask around it (radius=6 mm). Third, we extracted the mean value of beta within the mask from the subject and used it for further statistical analysis.

#### Psycho-Physiologic Interaction (PPI) model

We hypothesize that brain regions involved in computing have access to information about prior (*σ*_*π*_) and likelihood variability (*σ*_*L*_). Therefore, we predict that regions that represent *w*_*L*_ would exhibit stronger functional connectivity with regions that represent *σ*_*π*_ and *σ*_*L*_ at the time of prior and likelihood presentation. To test this prediction, we conducted a PPI analysis (Friston et al., 1997) and selected dACC that represented *σ*_*π*_ and *σ*_*L*_as the seed region. The GLM had the following regressors: (R1) The deconvolved and de-meaned seed time course (Gitelman, Penny, Ashburner, & Friston, 2003) (often referred to as the physiological regressor) and for each condition separately, the interaction between R1 and the indicator regressor at the time when prior and likelihood information were presented. In addition, all the regressors in GLM-2 were included in the model. All regressors were convolved with a canonical gamma hemodynamic response function. Temporal derivatives of each regressor were included in the model as regressors of no interest. The result of PPI analysis was corrected for multiple comparisons (z >2.3, family-wise error-corrected at p <0.05 using Gaussian random field theory)

## Acknowledgments

This work was supported by the Ministry of Science and Technology (MOST) in Taiwan (Grants MOST 104-2410-H-010-002-MY3 and 106-2420-H-010-003-to S.-W.W.) and by the Brain Research Center, National Yang-Ming University from The Featured Areas Research Center Program within the framework of the Higher Education Sprout Project by the Ministry of Education (MOE) in Taiwan. We acknowledge magnetic resonance imaging support from National Yang-Ming University, Taiwan, which is in part supported by the Ministry of Education plan for the top University.

## Notes

https://osf.io/ku97p/

